# Correlated stabilizing selection shapes the topology of gene regulatory networks

**DOI:** 10.1101/2022.08.29.505706

**Authors:** Apolline J. R. Petit, Jeremy Guez, Arnaud Le Rouzic

## Abstract

The evolution of gene expression is constrained by the topology of gene regulatory networks, as co-expressed genes are likely to have their expressions affected together by mutations. Conversely, co-expression can also be an advantage when genes are under joint selection. Here, we assessed theoretically whether correlated selection (selection for a combination of traits) was able to affect the pattern of correlated gene expressions and the underlying gene regulatory networks. We ran individual-based simulations, applying a stabilizing correlated fitness function to three genetic architectures: a quantitative genetics (multilinear) model featuring epistasis and pleiotropy, a quantitative genetics model where each genes has an independent mutational structure, and a gene regulatory model, mimicking the mechanisms of gene expression regulation. Simulations showed that correlated mutational effects evolved in the three genetic architectures as a response to correlated selection, but the response in gene networks was specific. The intensity of gene co-expression was mostly explained by the regulatory distance between genes (largest correlations being associated to genes directly interacting with each other), and the sign of co-expression was associated with the nature of the regulation (transcription activation or inhibition). These results concur to the idea that gene network topologies could partly reflects past correlated selection patterns on gene expression.

## Introduction

The development and physiology of living organisms are controlled by large and complex Gene Regulatory Networks (GRNs). The central role of GRNs is documented in all kinds of organisms, e.g. for the control of cell physiology in yeasts (Guelzim *et al*., 2002), heart development in humans (Olson, 2006), skeleton development in sea urchins (Shashikant *et al*., 2018), or flower development in angiosperms (Espinosa-Soto *et al*., 2004). The organization of these networks have long been of central interest for systems biologists, and it is now widely acknowledged that GRNs tend to follow general structural rules: for instance, they tend to be sparse, modular (Wagner *et al*., 2007; Espinosa-Soto, 2018), and scale-free (i.e., the number of connections per node follows a power law) (Babu *et al*., 2004; Ouma *et al*., 2018).

The reasons why real-life GRNs are organized in such a way are not completely clear (Espinosa-Soto, 2018; Taylor *et al*., 2022). Because the expression of genes affect phenotypic traits, and thus condition the individual fitness, gene expression levels are believed to be driven by natural selection, at least for a subset of genes. For instance, specific sets of genes have been shown to evolve in a direction consistent with prior knowledge in the wild (Philippe *et al*., 2007; Verta and Jones, 2019; Huang *et al*., 2021), or during experimental evolution (Philippe *et al*., 2007; Ghalambor *et al*., 2015; Jallet *et al*., 2020). In contrast, the structure of the network itself is less directly subject to natural selection. As multiple gene network topologies are capable of producing the same gene expression patterns, at least in theory (Wagner and Wright, 2007), the main evolutionary mode of network structure should follow non-adaptive processes, such as systems drift (Lynch, 2007), or mutation bias (Van Noort *et al*., 2004). Yet, a direct or indirect effect of selection on the evolution of network topology should not be excluded. For instance, it has been empirically established that the gene network structure may be deeply rewired during rapid evolutionary events, including domestication (Swanson-Wagner *et al*., 2012; Bellucci *et al*., 2014). Furthermore, the effect of indirect selection favoring evolvability (the propensity to produce mutant phenotypes with a good fitness) or robustness (the ability to buffer the effect of mutations) in gene networks remains a theoretical possibility (Wagner, 2008; Mayer and Hansen, 2017). Overall, there are only few theoretical predictions about how selection may affect the network topology, and about the possible role of adaptation in shaping GRN structure.

May the evolution of GRN topology be predicted from quantitative genetics theory? After all, gene expressions can be assimilated to quantitative traits, and the complex result of regulations can be described as epistasis (i.e., non-additive between genes) and pleiotropy (i.e., genes affect several traits) (Phillips, 2008; Fagny and Austerlitz, 2021). Evolutionary quantitative genetics provide a wide corpus of evolutionary models (e.g. Walsh and Lynch, 2018), including models designed to focus on the evolution of pleiotropy and modularity of quantitative characters (Sgrò and Hoffmann, 2004; Pavličev and Cheverud, 2015). With such theoretical tools, it has been showed that, if the genetic architecture is epistatic, pleiotropy could evolve in response to correlated stabilizing selection (Jones *et al*., 2014). Correlated selection, which corresponds to the selection of trait combinations (illustrated in Suppl. Fig. 1A), has been documented for various combinations of phenotypic characters (Sinervo and Svensson, 2002), but its consequences on the structure of genetic architectures is not well understood (Uller *et al*., 2018; Svensson and Berger, 2019; Svensson *et al*., 2021). The evolutionary mechanism involved in the evolution of pleiotropy relies on the fact that the genetic load of new mutations is minimized when the mutational correlation matches the direction of the fitness function (Suppl. Fig. 1B). In other terms, the effect of mutations are expected to evolve to promote trait combinations favored by selection (Jones *et al*., 2007). As the mutational effects are a direct consequence of the genetic structure, the simulations by Jones *et al*., 2014 thus formalises the hypothesis that correlated selection could favor gene network topologies promoting the co-expression of co-selected genes. Yet, this important result from evolutionary quantitative genetics may not be straightforward to translate towards systems biology, as the genetic architecture in Jones *et al*., 2014 was based on a bivariate multilinear model (Hansen and Wagner, 2001), featuring unconstrained and isotropic pleiotropic epistasis (i.e., any gene have the potential to modify the pleiotropy of any other gene). In contrast, the epistatic and pleiotropic effects in GRNs are largely constrained and biased by the topology of gene networks (Sorrells *et al*., 2015; Nghe *et al*., 2018).

Here, we intend to understand the propensity of correlated stabilizing selection to shape the structure of gene networks. We will use the theoretical framework proposed by Wagner, 1994, 1996 to implement a simple gene regulatory network model as a genotype-phenotype map. We will monitor the evolution of pleiotropy among gene expressions in individual-based simulations. This framework is well-suited for being coupled with simulations, as the genotype (the set of regulations between genes) and the phenotype (gene expressions) are explicit and clearly separated. We will address the evolution of gene co-expression at two levels: (i) at the gene expression level, can gene networks evolve to optimize mutational correlation in regard to correlated selection ? (ii) at the network level, what is the effect of correlated selection on network structure and topology? The evolution of co-expression in the GRN model will be compared to the evolution of pleiotropy in two quantitative genetics models: the bivariate multilinear model (Hansen and Wagner, 2001; Jones *et al*., 2014) and the gene pleiotropy model (Lande, 1980).

## Material and Methods

Our purpose is to measure the evolutionary changes in the properties of the genetic architectures when submitted to correlated selection, with a particular focus on the propensity of mutations to induce pleiotropic (correlated) effects on co-selected phenotypic traits. The influence of the nature of the genotype-phenotype relationship will be addressed by considering three genotype-phenotype models, explored by individual-based simulations.

### Measurement of pleiotropy *via* the mutational covariance matrix

In multivariate quantitative genetics models, the response to directional selection in a complex phenotypic space can be predicted from the structure of the (additive) genetic covariances (the **G**-matrix) in the population (Lande and Arnold, 1983; Blows, 2007). Genetic covariances result from both linkage disequilibrium (LD), the statistical association of alleles at different loci, and pleiotropy. LD is reversible, it can be affected by genetic drift, recombination rate, recent directional or stabilizing selection, and gene flow. In contrast, pleiotropy reflects the properties of the genetic architecture of the traits, and is generally considered as a non-evolvable constraint when studying the adaptation of quantitative traits (e.g. Jones *et al*., 2003; Chantepie and Chevin, 2020).

Here, our objective is to study the long-term evolution of pleiotropy as a consequence of selection for trait combinations. Pleiotropy can be formally measured as the propensity of mutations to affect two or more traits together. The distribution of the multivariate effects of mutations can be summarized by the matrix **M**, which diagonal and off-diagonal elements stand for mutational variances and covariances respectively. Most of the following results will focus on two traits, named *a* and *b*; the corresponding **M** being:

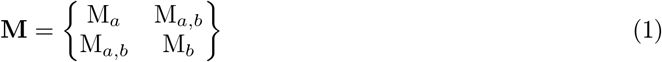

where M_*a*_ and M_*b*_ are the mutational variances of traits *a* and *b*, respectively, and M_*b,a*_ = M_*a,b*_ is the mutational covariance between traits *a* and *b*.

Two-dimensional covariance matrices can be conveniently represented graphically as ellipses (Cheverud, 1984; Jones *et al*., 2014), sometimes assimilated to the corresponding 95% confidence interval of a multivariate Gaussian distribution. We will extract two geometrical properties from these matrices, the direction (angle) between its main eigenvector and the first trait, measuring the main mutational direction *α*(**M**), and the ellipse eccentricity *e*(**M**), measuring the strength of pleiotropy from 0 to 1.

The calculation of the mutational direction is detailed in the Supplementary Methods section; the eccentricity of the **M** matrix was computed as 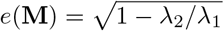, where *λ*_*i*_ stands for the *i*^th^ eigenvalue of the matrix **M**. Mutational correlation *r*(**M**) between genes *a* and *b* were calculated from **M** matrices using the standard formula 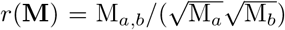. The relationship between direction, eccentricity, and correlation is illustrated in Figure 1A.

**Figure 1.**
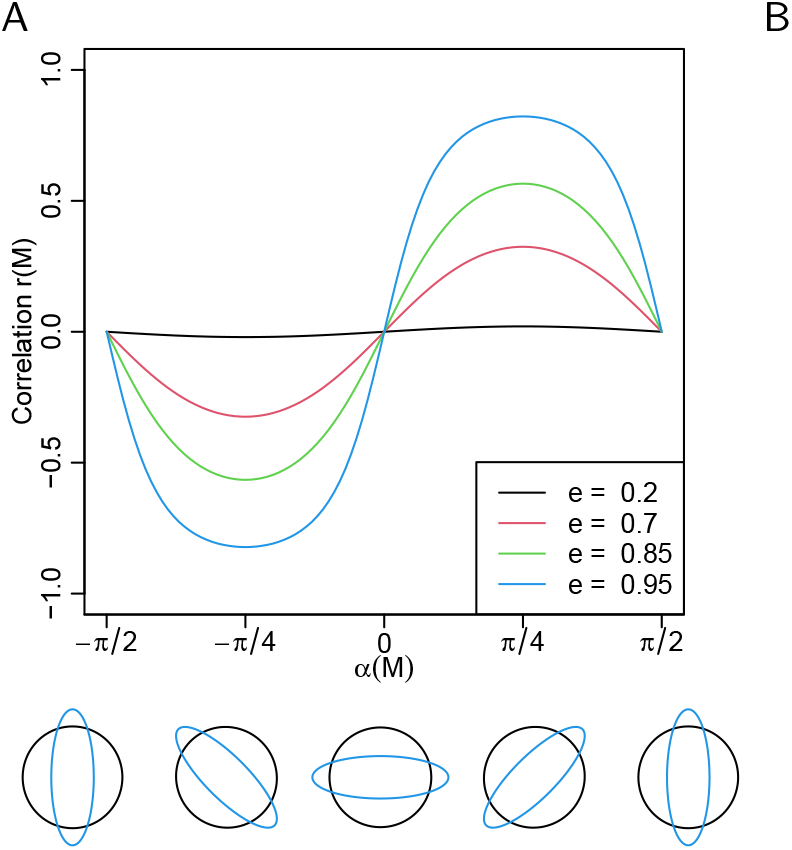
A: The relationship between the eccentricity (*e*), direction (*α*) and correlation (*r*) of a variance-covariance matrix **M**. The geometry of matrices with different directions (**M** = − *π*/2, − *π*/4, 0, *π*/4, − *π*/2) and eccentricities (black: *e* = 0.2, blue: *e* = 0.95) is represented below the x-axis. B: Diagram representing our gene regulatory network design. *a* and *b* are the focal genes (the genes which expression is under correlated selection). *c* and *d* are genes selected to be activated (optimal phenotype at **Θ**_*i*_ = 0.5, corresponding to an optimal expression ≃ 0.62, slighly above the basal expression *κ* = 0.5), independently from each other. *e* and *f* are free to evolve without affecting the fitness directly, and can thus act as transcription factors.

While the genetic covariance matrix **G** is a population property, the mutational matrix **M** is a property of a genotype. **M**_*i*_ was thus estimated for every individual *i* of the population, and variances and covariances were averaged out to get the population **M**. Thirty independent simulation replicates were run, and some figures report average values. Average correlations 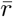 and eccentricities 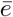 were computed as arithmetic means, while the mean direction 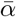 over *R* replicates was obtained as a circular mean restricted to the interval (−*π*/2, *π*/2) (detailed in the Supplementary Methods).

### Selection

Relative fitness was determined by a multivariate stabilizing bell-shaped fitness function (Lande, 1980):

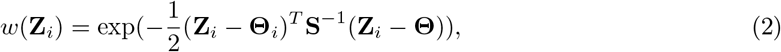

where **Z**_*i*_ is the vector of phenotypes for individual *i*, **Θ**_*i*_ is the optimal phenotype for trait *i* (by default, **Θ**_*i*_ = 0 unless specified otherwise), and **S** is the covariance matrix of the fitness function. The trace of the matrix **S** (the sum of the diagonal elements) represents the width of the fitness function (the larger the coefficients of **S**, the weaker the selection). The fitness function was parameterized so that the maximum relative fitness was *w*(**Θ**) = 1.

For simplicity, the number of phenotypic traits on which correlated selection was applied was reduced to two traits in most simulations. As a consequence, the fitness function was specified by five parameters: two parameters for the phenotypic optima, and three (co)variance parameters for the 2 × 2 matrix **S** (the strength of selection on traits 1 and 2, and the selection correlation *r*(**S**)). The main direction of selection *α*(**S**), the eccentricity *e*(**S**) and the correlation *r*(**S**) of the fitness function have the same meaning as for the mutation covariance matrix.

### Genotype-Phenotype models

To simulate the evolution of populations and their **M** matrices, we used three models implementing different genotype-phenotype mapping. The first model is a gene regulatory network (GRN) model, in which the genotype represents regulations between transcription factors, and the phenotype is the expression of the network genes at equilibrium. The second model is a bivariate version of the multilinear model (Hansen and Wagner, 2001; Jones *et al*., 2014), which extends the classical additive model with epistatic and pleiotropic interactions. The third model (that we called the Gene Pleiotropy model, GP) is based on an implementation of Fisher, 1930‘s geometric model in which every additive locus has its own pleiotropic pattern (Lande, 1980).

#### Gene regulatory network model

We used a regulatory gene network model inspired from Wagner (Wagner 1994, 1996), which is a common abstraction of the transcription regulation process as it is a dynamic model with discrete time steps (Bergman and Siegal 2003; Azevedo *et al*. 2006; Leclerc 2008; Rhoné and Austerlitz 2011; Rünneburger and Le Rouzic 2016; Espinosa-Soto 2016). The structure of the regulation network among *n* genes is stored in a *n* × *n* matrix **W**, corresponding to cis-regulations among transcription factors. Element *W*_*ij*_ corresponds to the effect of the product of gene *j* on the expression of gene *i*. Inhibiting regulations are negative values and activating regulations are positive values. Zero indicates the absence of direct regulation. Each gene of the network is susceptible to act as a transcription factor and to modify the expression of other genes; there was no self-regulation (*W*_*ii*_ = 0).

Gene expression was computed dynamically for 24 time steps, which happens to be enough to reach equilibrium in our simulations (Suppl. Fig. 5). Initial gene expressions were set to their basal level (expression in absence of regulation) *κ* = 0.5, intermediate between full inhibition and full activation. Gene expressions were dynamically updated as a function of the concentration of the other genes of the network:

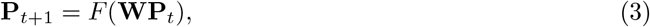

where **P** is a vector of *n* quantitative gene expressions, scaled between 0 (no expression) to 1 (maximal expression), by a sigmoid function *F* (*x*_1_, …, *x*_*n*_) = (*f* (*x*_1_), …, *f* (*x*_*n*_)). This scaling function was *f* (*x*) = 1/(1 + *e*^−4*x*^) for the default basal expression *κ* = 0.5 (see supplementary methods for *κ* ≠ 0.5).

The phenotype **Z** corresponding to a genotype **W** was computed from the average expression of the two first genes of the network (hereafter called “*a*” and “*b*”) for the 4 last time steps 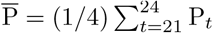, as 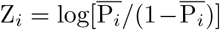, rescaled over (−∞, +∞) to be directly comparable with the multilinear model; *Z*_*i*_ < 0 corresponds to underexpression, *Z*_*i*_ > 0 to overexpression, and the phenotypic value *Z*_*i*_ = 0 to an expression intermediate between the minimum and maximum.

Network topology and the corresponding gene expression evolved because the strength of regulation *W*_*ij*_ can change by mutation (except for self-regulation *W*_*ii*_, which was set to a constant 0). The mutation rate per individual was *μ*, and each gene had the same probability *μ/n* to be affected by a mutation. Mutations changed a single random element of the mutated gene by a Gaussian deviation of variance *σ*^2^*m* (see Table 2 for parameter values).

In order to facilitate the evolution of diverse regulatory motifs in the network (involving more than the two target genes), two genes (*c* and *d*) were considered as “transcription factors”, and selected to be up-regulated by including them in the fitness function (equation 2), with an optimum *θ*_*c*_ = *θ*_*d*_ = 0.5 (corresponding to an optimal expression of *P*_*c*_ = *P*_*d*_ = 0.62) and a selection strength of *S*_*c,c*_ = *S*_*d,d*_ = 10, selection being uncorrelated (*S*_*c,i* ≠ c_ = 0) (see Figure 1B). In addition to the selection on the phenotype, unstable networks were penalized (considering unstable networks as unviable is common in the literature, see e.g. Siegal and Bergman 2002). In practice, the individual fitness *w* was multiplied by a factor 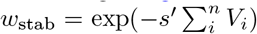 where *s*^*′*^ quantifies the selection against unstable gene expression, and 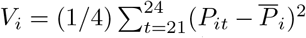 is the variance in the expression of gene *i* during the last 4 steps of the network dynamics (more details in Supplementary methods). We set *s*^*′*^ = 46, 000, as in Rünneburger and Le Rouzic 2016, which was a large penalty; in practice, unstable networks were thus strongly selected against and these genotypes were absent from the simulations except for rare spontaneous mutants.

#### Multilinear model

The multilinear model was originally developed by Hansen and Wagner, 2001. Although provided as a multivariate model in its original description, it has been extensively used in its univariate form in the quantitative genetics literature (Hermisson *et al*., 2003; Carter *et al*., 2005; Jones *et al*., 2007; Le Rouzic *et al*., 2013), but more rarely in its multivariate implementation (Jones *et al*., 2014).

The multilinear model is built as an extension of the additive model, by adding epistatic terms proportional to the product of the additive effects across genes. Restricting the model to second-order epistasis (interactions between pairs of genes), the phenotypic value *Z*_*m*_ of a trait *m* (among *K* traits) is:

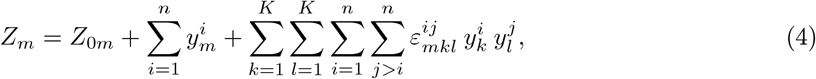

in which 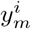 is the effect of the genotype at gene *i* on the phenotypic trait *m* measured in an arbitrary reference genotype where all 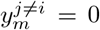. *Z*_0*m*_ is the phenotypic reference, i.e., the phenotypic value corresponding to an arbitrary reference genotype for which all 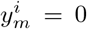. For every combination of traits, the epistatic coefficient *ε*^*ij*^ quantifies the directional epistasis between genes *i* and *j*. The coefficients *ε*_*mmm*_ describe “classical” epistasis, i.e., the interaction of allelic effects 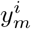 and 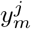 on trait *m*. In contrast, *ε*_*mkl*_ with *k* and/or *l* different from *m*, correspond to interactions involving pleiotropy, i.e., how trait *m* is influenced by the interaction between the effects of alleles on traits *k* and *l*. When all *ε*^*ij*^ = 0, this model collapses towards an additive model. When *ε*_*mkl*_ ≠ 0, pleiotropy can evolve (traits can become more or less dependent). In total, there are *K*^3^ combinations of *K* traits, and for *n* genes, *n*(*n* − 1)/2 independent epistatic coefficients (because *j > i*) for each combination of traits.

In the multilinear model, evolution occurs because 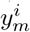 can change. Mutations affect genes independently, and a mutation at gene *i* affects all traits at once (the effect of mutations being independently drawn in Gaussian distributions of variance 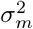). In contrast, the *ε* coefficients (2^3^ × 6 × 5/2 = 120 in the default setting) coud not evolve. They were drawn in a Gaussian distribution *ε* ∼ 𝒩 (0, 1) at the beginning of each simulation run and kept constant throughout generations, as in Jones *et al*., 2014.

#### Gene Pleiotropy model

We also considered a model in which gene contributions were additive (i.e., pleiotropy was not modeled as epistasis), but each locus had its own pleiotropic structure (termed “orientation heterogeneity” in Chevin *et al*., 2010). This setting is inspired from Lande, 1980 and is regularly used to study the evolution of modularity (Chevin *et al*., 2010).

In practice, every gene *i* was featured by its own mutational matrix **M**_*i*_ = *μ*_*i*_**C**_*i*_, where the covariance matrix **C**_*i*_ quantifies the pleiotropy at gene *i*. The covariance matrix **C**_*i*_ was constant, but the mutation rate *μ*_*i*_ was evolvable, opening the possibility for the gene to increase or decrease its overall contribution to the mutational properties of the genotype. Gene-specific covariance matrices **C**_*i*_ were computed in order to cover equally-spread angles between −*π*/2 and *π*/2, with a strong eccentricity (*e*(**C**_*i*_) = 0.9). The genotype was encoded in the same way as in the multilinear model, 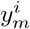 being the additive effect of gene *i* on trait *m*, and the genotype-phenotype map was additive 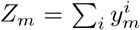. There are two kinds of mutations: “regular” mutations affecting the traits (“trait mutations”), and mutations affecting the gene mutation rate (“rate mutations”). Trait mutations occurred with a rate *μμ*_*i*_/ ∑_*j*_ *μ*_*j*_ at gene *i*; they affect all traits at once, and mutational effects were correlated according to the covariance matrix **C**_*i*_. Mutation rates were normalized so that the mutation rate per individual and per generation is *μ*, as in the other models (the mutation rate of genes evolved relative to each other, but the total mutation rate remained constant). Rate mutations occurred with a rate *μ*\* per genotype and per generation (for convenience, *μ*\* = *μ*), and may affect all loci with the same probability. Their effect was Gaussian on the multiplicative scale (the mutation rate after mutation was 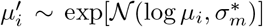, and the effect of rate mutation was fixed to 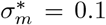 (in average, a rate mutation changed the mutation rate by ≃ 8.3%).

The similarities are differences among the three models are summarized in Table 1.

**Table 1:**
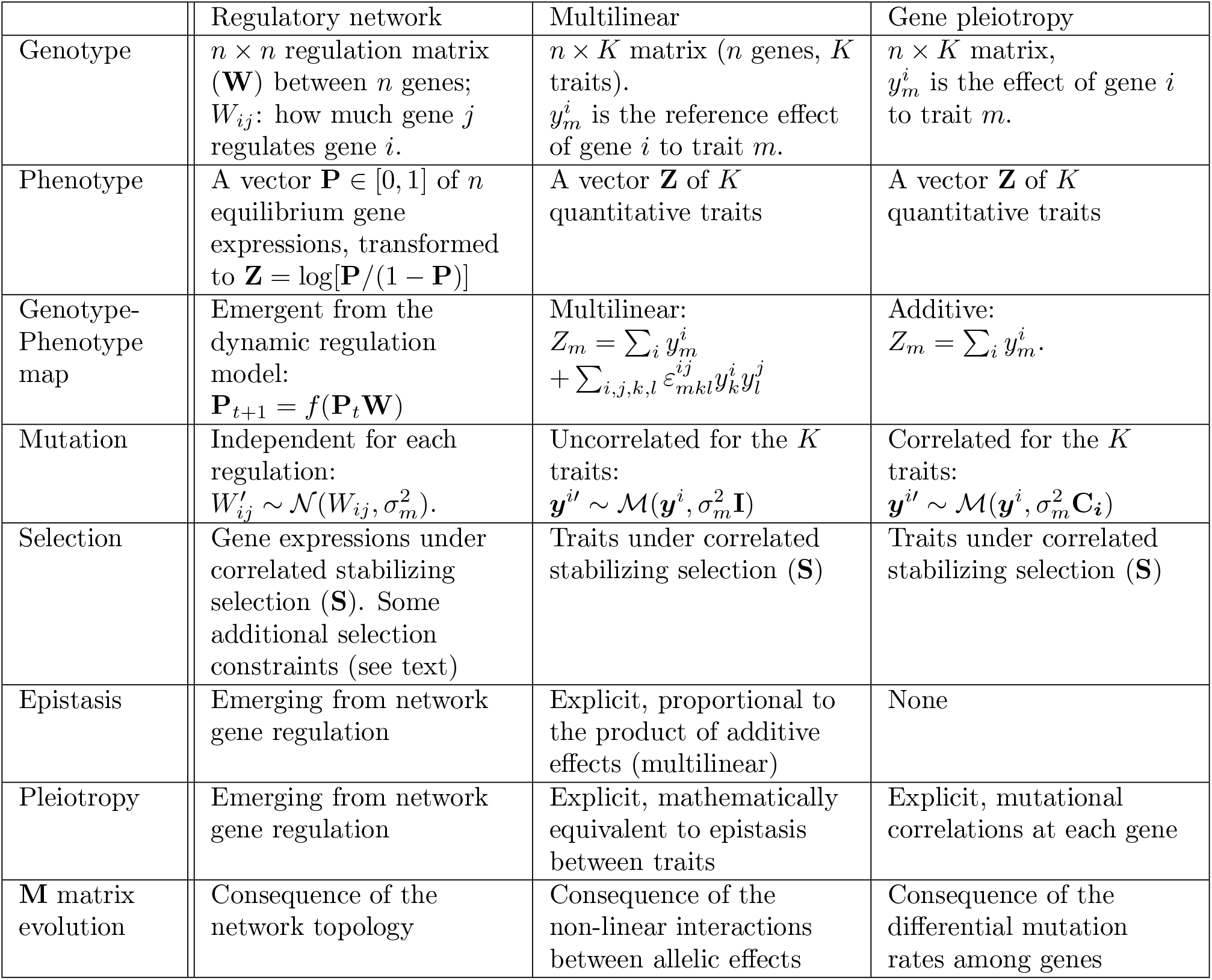
Comparative table for the three models. 𝒩(*μ, σ*^2^): Normal distribution of mean *μ* and variance *σ*^2^; ℳ (***μ*, Σ**): Multivariate normal distribution of means ***μ*** and covariances **Σ. I** stands for the identity matrix of the adequate dimension. Other symbols are described in the text.

**Table 2:** Default parameters for the three models.

| Parameter | GRN | Multilinear | Gene pleiotropy |
| --- | --- | --- | --- |
| Generations | 10000 | 10000 | 10000 |
| Population size $N$ | 5000 | 5000 | 5000 |
| Genes $n$ | 6 | 6 | 6 |
| Correlationally selected traits | 2 | 2 | 2 |
| Haplotype mutation rate $\mu$ | 0.1 | 0.1 | 0.1 |
| Mutation effect $\sigma_m$ | 0.1 | 0.0369 | 0.0369 |
| Optimum phenotype | 0 | 0 | 0 |
| **S** matrix size (trace) | 10 | 10 | 10 |
| **S** matrix eccentricity | 0.94 | 0.94 | 0.94 |

### Simulation model

All data presented in this article have been generated by computer simulation of evolving populations with the C++ program Simevolv (Rünneburger and Le Rouzic 2016: https://github.com/lerouzic/simevolv.git). The analysis scripts have been written in R and are available at https://github. com/apetit8/Mmatrix_paper.git.

#### Reproduction

Simulations followed a traditional Wright-Fisher framework. Populations consisted in *N* haploid, sexually-reproducing hermaphrodite individuals. The genotype was encoded as *n* freely recombining genes, the (multivariate) phenotype being computed from the genotype according to one of the genotype-phenotype models described above. Generations were non-overlapping; for each offspring, two parents were picked with a probability proportional to their relative fitness, and the two *n*-gene haploid gametes were recombined to form a new haploid genotype. Populations evolved during 10,000 generations and were submitted to genetic drift, selection, and mutations.

#### Mutations

Mutations affect the genotype immediately after recombination, before the computation of the phenotype of individuals. Mutations occurred with a rate *μ* per gamete, and affect random genes as described above. Mutational effects were cumulative, the new allelic value was drawn in Gaussian distributions centered on the former values.

#### Model output

The simulation software reports the means, variances and co-variances of the population phenotypes and genotypes at regular time points. In addition, the population average mutation co-variance matrix **M** was estimated in the following way: 6 mutations were simulated for each of the *N* = 5, 000 individuals *i*, leading to 5, 000 covariance matrices that were averaged out and multiplied by the mutation rate *μ*.

### Simulation parameters

Default simulation parameters were set as displayed in Table 2. For the multilinear model, the epistasis parameters were inspired from Jones et al., 2014 (Jones *et al*., 2014). Parameters for the three models were adjusted to produce M matrices of similar sizes.

All simulations starts with genotypic values (*y*_*i*_ in the multilinear and GP models, *W*_*ij*_ in the gene network model) set at 0, unless specified otherwise. In some simulations, the initial gene network topology was manipulated (positive or negative initial correlation) by setting some initial regulations (*W*_*ab*_ and *W*_*ba*_) with positive (+0.5), negative (− 0.5), or null (0) values. The corresponding slots of the **W** matrix (*W*_*ab*_ and *W*_*ba*_) were not evolvable and remained to their initial values, while the rest of the network was free to evolve.

The bivariate stabilizing selection (**S** variance matrix) was parameterized in each simulation run by setting the angle of the major axis (between − *π*/2 and *π*/2); the matrix size tr(**S**) and eccentricity remained constant (see Table 2 and orange ellipses in Figure 2).

**Figure 2.**
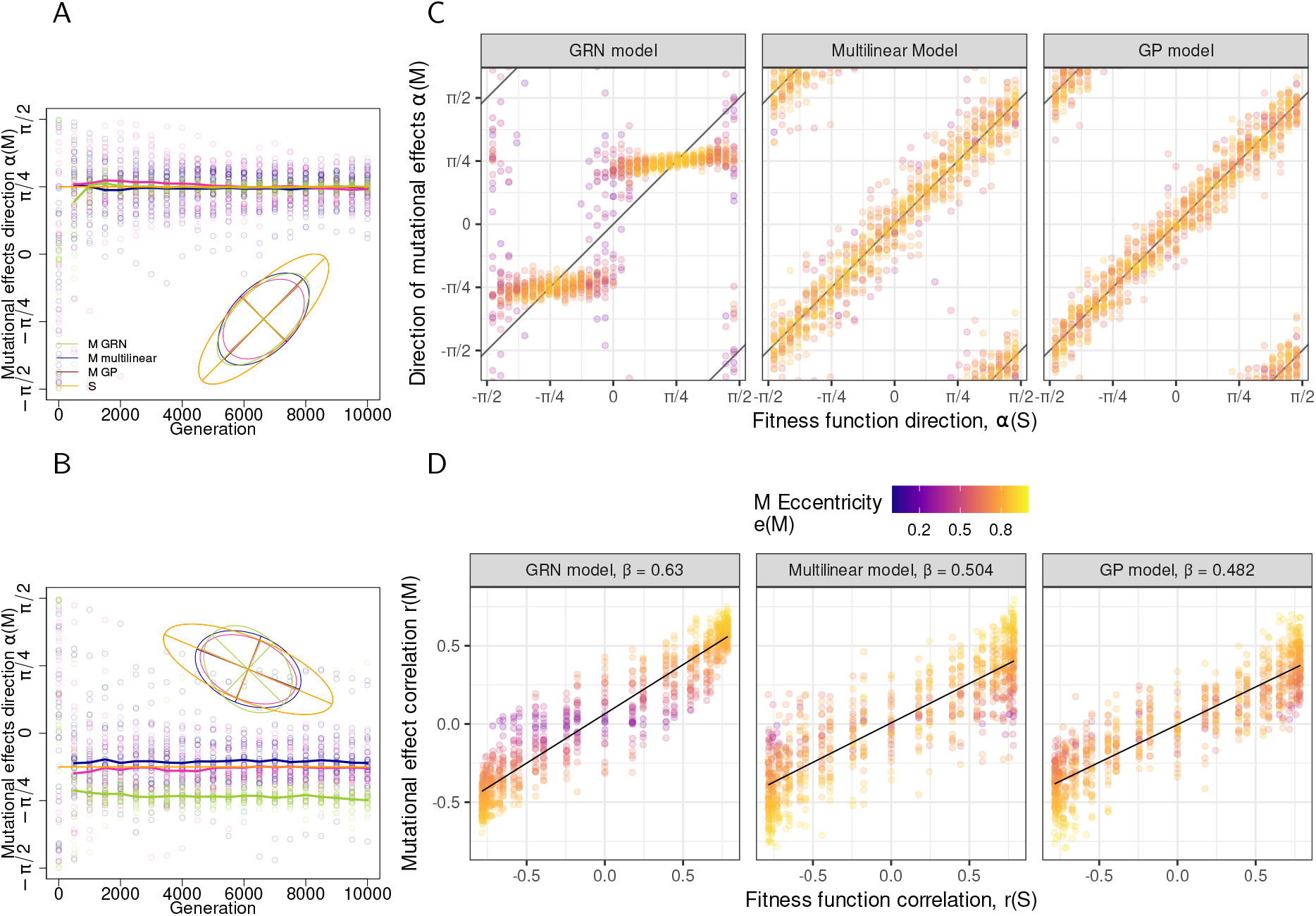
A, B: Evolution of the angle of the main axis of the mutation matrix (*α*(**M**)) along generations. Orange ellipses illustrate the fitness function (scaled × 0.025), which direction was *α*(**S**) = +*π*/4 (panel A), and *α*(**S**) = − *π*/8 (panel B). Dots represent 30 simulation replicates, plain lines stand for circular means 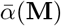. Ellipses are the geometric representation of **M** matrices at the last generation (10,000), averaged over the 30 replicates. C: *α*(**M**) as a function of *α*(**S**) for the three models. Dots represents the direction of the main axis of the **M** matrix. D: *r*(**M**) as a function of *r*(**S**) for the three models. For C and D: Data obtained after 10,000 generations in 30 simulation replicates for 31 values of *α*(**S**) regularly spaced between − *π*/2 and *π*/2; the color scale encodes the eccentricity of **M**. *β* is the linear regression coefficient between *r*(**M**) and *r*(**S**).

## Results

### Mutational correlations can evolve in all models

We compared the evolution of simulated populations based on three genetic architectures: a gene regulatory network architecture (GRN model), considering gene expression levels as phenotypic traits, quantitative traits controlled by a multilinear genetic architecture (as in Jones *et al*., 2014), additive traits controlled by several genes displaying different peiotropic patterns (Lande, 1980) (GP model). Two traits (two gene expressions for the GRN model) were submitted to correlated stabilizing selection, the fitness function being defined by the direction *α*(**S**) of the optimal trait combination. Our expectation was that pleiotropy (measured as the shape and direction of the mutational covariance matrix **M**) should evolve in order to match the direction of the fitness function.

In the multilinear and GP models simulations, the alignment between the main axis of the mutational matrix **M** and the direction of the correlated fitness function was convincing after less than 500 generations (Figure 2A, B). This result confirmed the conclusions from Jones *et al*., 2014, based on the multilinear model. In contrast, our gene network model did not always evolved towards the best alignment, even after 10,000 generations (e.g. in Figure 2B): the sign of mutational correlations matched the sign of fitness correlations, but there was a discrepancy at equilibrium.

The different nature of the response to correlated selection in the three models is illustrated in Figure 2C. For both the multilinear and GP models, the response to the direction of the fitness function *α*(**S**) was homogeneous in all directions, and the shape (eccentricity) of the **M** matrix did not depend on *α*(**S**). In contrast, with the GRN model, although both the direction and eccentricity of **M** evolved, pleiotropy evolved along preferential directions: *α*(**M**) did match the sign of *α*(**S**), but not the precise direction of the fitness function. Intermediate angles (*π*/4: both gene expressions affected equally by mutations, and − *π*/4: opposite effects on both genes) were frequently observed, and mutational independence (*α*(**M**) = ±*π*/2 or 0) was difficult to achieve. In the GRN model, evolving different mutational effects for both selected traits was more difficult, leading to frequent round (weak eccentricity) **M** matrices. This appears to reflect a property of gene network architectures, as GRNs tend to evolve towards this pattern even when starting from a better alignment (Suppl. Fig. 3).

While pleiotropy (and absence of pleiotropy) could not evolve in the GRNs as freely as in the other models, GRN models displayed the best response of mutational effect correlation *r*(**M**) to the fitness correlation *r*(**S**): *β*_GRN_ (linear regression coefficient) of 0.63, against *β*_GP_ = 0.48 and *β*_multilin_ = 0.50 (Figure 2D). The constrains on the evolution of pleiotropy did not translate into the genetic covariance matrix **G**, which was aligned on selection for all models (Suppl. Fig. 4) due to the contribution of linkage disequilibrium. Overall, all three models evolve under correlated selection through different strategies : the direction of the **M** matrix tends to evolve quantitatively in the GP and multilinear models, while the response of GRNs is rather discrete (positive, negative, or no pleiotropy).

### Mutational correlation is determined by local regulatory motifs

In the previous section, we used quantitative genetics tools to describe the structure of mutational correlations among traits and its evolution. Here we aim at deciphering the changes in the regulatory motifs that underlie the evolution of co-expression in gene networks.

We measured the correlation between each of the 30 network regulations **W**_*ij*_ and the quantitative descriptors of **M** (*α*(**M**), *e*(**M**) and *r*(**M**)) (Figure 3). The regulations affecting **M** the most were the direct regulations between target genes *a* and *b*. In contrast, regulations between the rest of the network (especially the overexpressed transcription factors *c* and *d*) towards the focal genes *a* and *b* decreased pleiotropy. The other regulations did not affect the direction or eccentricity of the mutational matrix, strongly suggesting that the co-expression between two genes is determined by the local regulatory motif.

**Figure 3.**
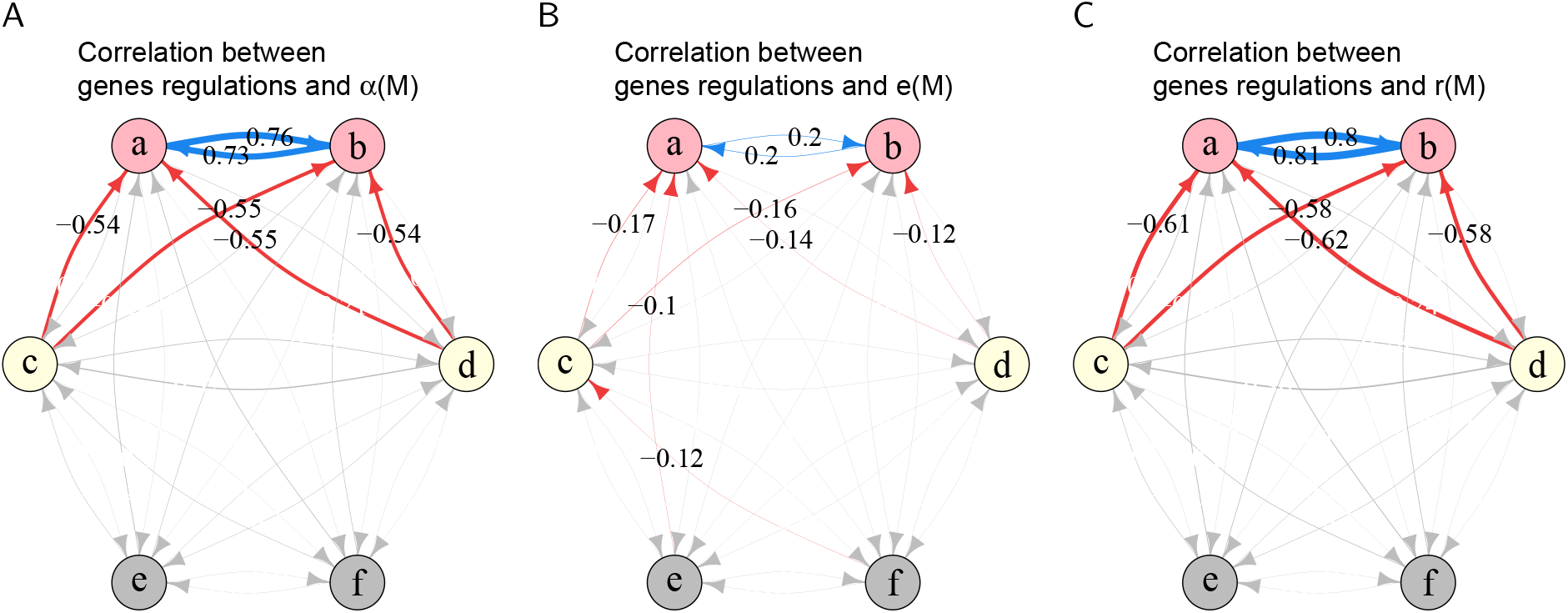
Network graph (off-diagonal elements of the regulatory matrix **W**) with edges width proportional to the correlation between genes regulations **W**_*ij*_ and A: the mutational matrix direction *α*(**M**), B: the mutational matrix eccentricity *e*(**M**), and C: the mutational correlation *r*(**M**). Positive correlation are blue, negative correlations are red. Correlations inferior at 0.4 for A and C and inferior at 0.1 for B are represented in grey. Data used to calculate correlations are the one represented in Figure 2D (930 networks in total, corresponding to 31 regularly distributed angles for the correlated fitness function, with 30 simulation replicates for each angle).

The influence of direct regulations was further assessed by forcing their value to positive (activation), negative (repression), or zero (no regulation allowed). Fixing regulations between the focal genes prevented the evolution of the direction of the mutational matrix (Figure 4A and B), and *α*(**M**) was constrained by the sign of the direct regulation (mutually activating genes were always positively coexpressed, mutually inhibiting genes were always negatively co-expressed). When direct regulations were prevented, co-expression could evolve, but to a lesser extent (Figure 4C). Direct regulations are thus the main contributor to the evolution of the pleiotropy in gene networks.

**Figure 4.**
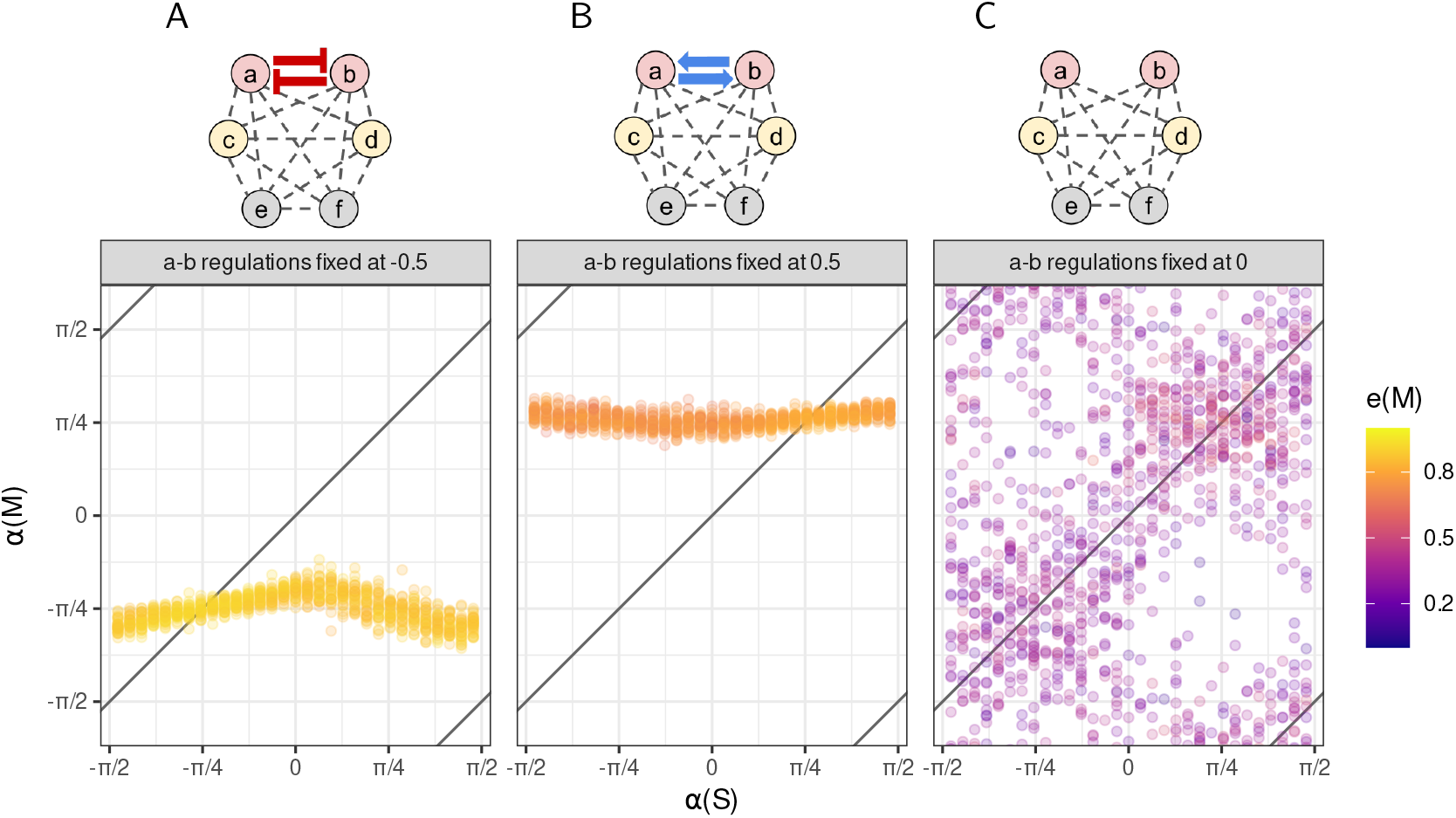
Direction of mutational matrices (*α*(**M**)) after 10,000 generations of evolution as a function of the direction of the correlated fitness function (*α*(**S**)). The representation is the same as in Figure 2D and E. A: fixed negative (inhibition) regulation between genes *a* and *b*, B: fixed positive (activation) regulation, C: no regulation. The corresponding networks are represented above the scatterplots (red arrows: constant inhibition, blue arrows: constant activation, grey hyphenated connections: evolvable regulations; whether or not such regulations evolved in the simulations differed among replicates). Other conditions were the same as in Figure 2E.

We manipulated the network topology to increase further the network distance between focal genes and assessed the effect of network distance on mutational correlations (Figure 5). While direct regulations between two genes allowed for the evolution of mutational correlations ranging from − 0.6 to +0.7, correlation intensity decreased with the network distance, as it barely spanned ± 0.2 with one intermediate gene, and ±0.05 with two intermediate genes. No mutational correlation was detected with more than two intermediate genes. Similar simulations with larger networks displayed the same trend (Suppl. Fig. 5).

**Figure 5.**
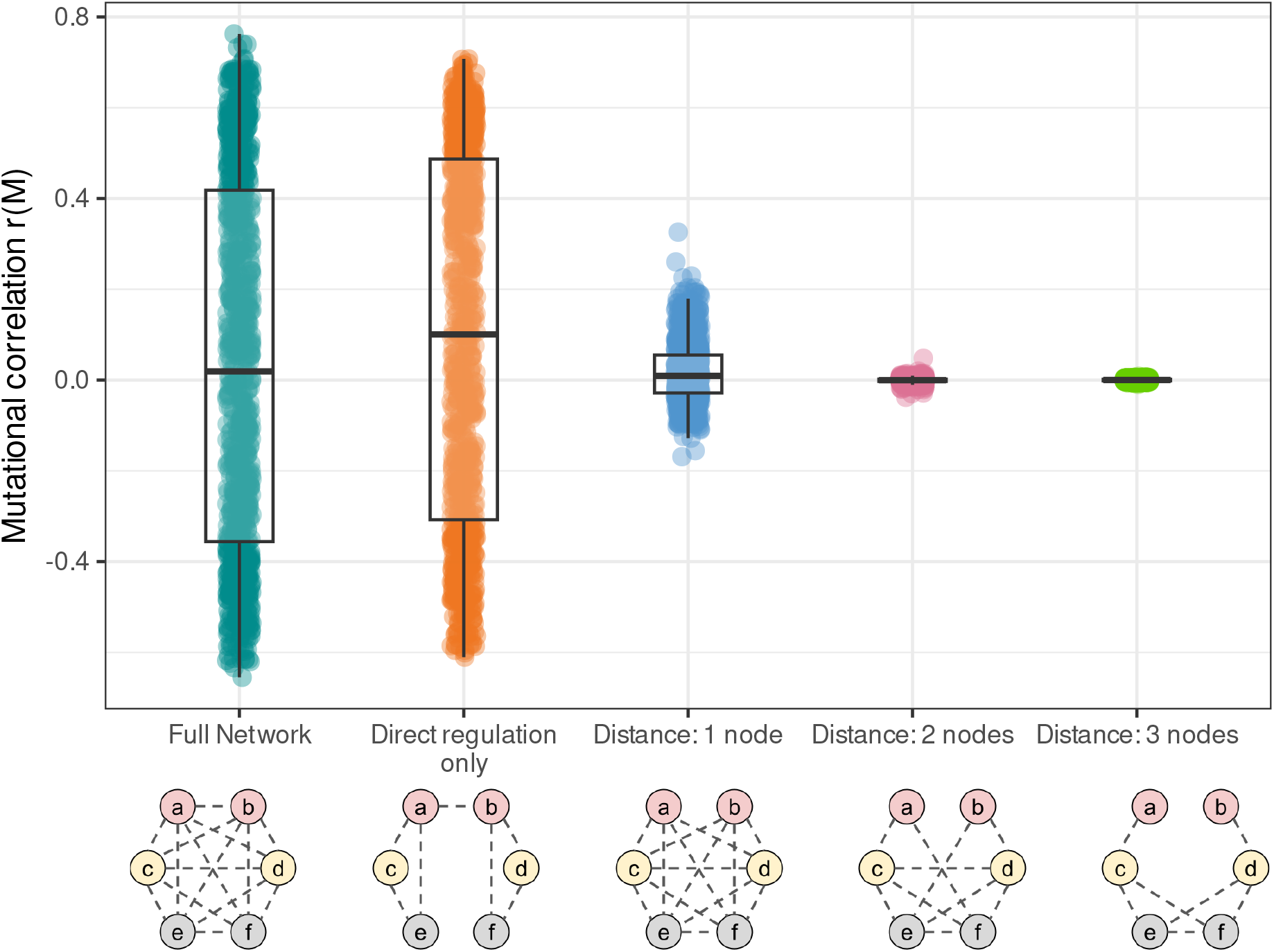
Distribution of mutational correlations *r*(**M**) in networks of different topologies, illustrated under the x-axis. All networks have evolved in the same set of conditions as in Figure 4, i.e., 31 correlated fitness functions oriented in various directions. Genes colored in pink (*a* and *b*) are the genes under correlated selection, yellow genes (transcription factors *c* and *d*) are under non-correlated selection, and grey genes are not selected (sans color code as in Figures 1 and 4). Regulatory connections indicate which regulations were possible; whether or not such regulations evolved in the simulations differed among replicates.

### Correlated selection can shape gene networks at a large scale

So far, we assessed whether the fitness correlation between two genes only could shape the local topology of the gene network, which was arguably a favorable scenario that maximizes the chances for the genetic architecture to respond. Realistic selection pressures are probably more complex, involving many genes and a complex pattern of fitness correlations among them. To check whether the observed pattern was maintained at a larger scale, we simulated the evolution large GRNs in which all genes were under stabilizing selection (diagonal elements of the matrix **S** were set to 10), while all pairwise correlations were drawn randomly. Although the association between the fitness correlation and the mutational correlation weakened with the number of genes, the response was still observable with up to 30 genes (Figure 6). This confirms that network topology can be shaped by selection at a large scale, and that our results are not an artifact of focusing on small network motifs.

**Figure 6.**
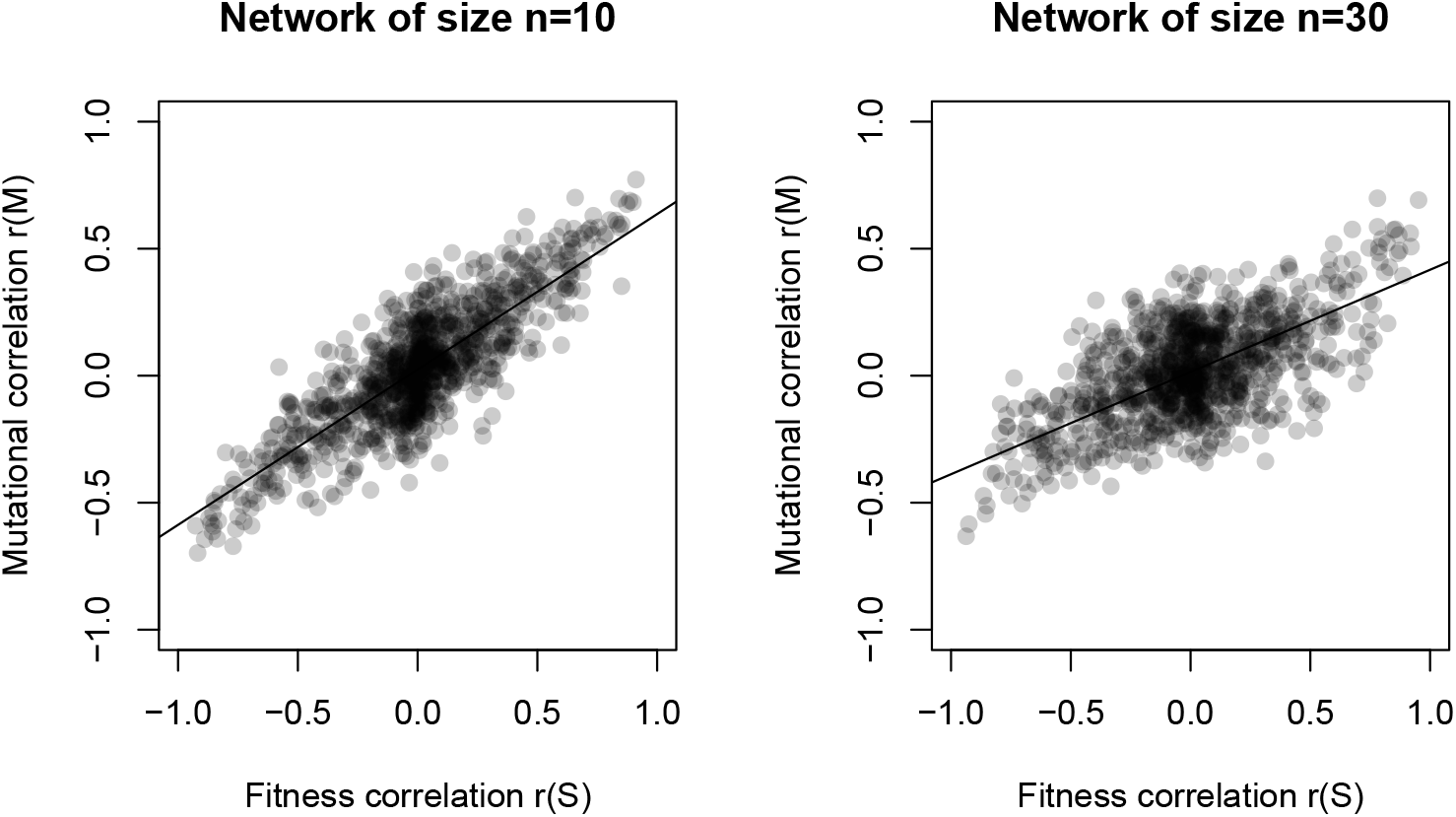
Mutational correlation after 20000 generations of evolution, as a function of the fitness correlation in networks of various sizes (*n* = 10 and *n* = 30). Each pair of genes was under correlated selection, correlations ranging from −0.9 to 0.9. To ensure that the correlation matrix was positive definite, selection correlations were the non-diagonal elements of the *n* × *n* matrix ***uu***^T^, where *u* was drawn in a uniform (−1; 1) distribution. The mutation rate was scaled across simulations to ensure a constant mutation rate by gene.

The effect of network size (*n*), as well as five other parameters (population size *N*, mutation size *σ*_*m*_, strength of selection, and basal expression *κ*) have been explored in Suppl. Fig. 6. Our main result (the mutational structure responds to correlated selection) appeared to be robust to parameter changes, and may only be affected by extreme parameter values. The default parameter set was not necessarily optimal, as larger co-expression could evolve in large populations (*N* = 10, 000).

## Discussion

Constant stabilizing selection is often thought to promote stasis, and thus prevent the evolution of traits, but it is also suspected to modify the structure of genetic architectures, in particular through the minimization of the fitness load due to environmental disturbances and mutations (Wagner *et al*., 1997). When considering phenotypic traits independently, stabilizing selection promotes genetic backgrounds that reduce mutational effects (Rice, 2002; Hermisson *et al*., 2003). Simulations based in gene network models have reproduced this predictions, and have associated the decrease in the effects of mutations with systematic changes at the network level, including feedback loops, global network size and properties, and redundancy (Masel and Siegal, 2009; Payne and Wagner, 2015; Rünneburger and Le Rouzic, 2016).

Quantitative genetics theory predicts that mutational correlation between traits (pleiotropy) can evolve as the result of correlated stabilizing selection (Cheverud, 1984; Jones *et al*., 2014). Yet, this theoretical evolutionary force is weak and indirect, and could easily be overwhelmed by genetic drift, correlations with phenotypic trait values, and mutation bias. We simulated the evolution of mutational correlations in three genetic architectures to assess whether the evolution of pleiotropy depends on the mechanisms underlying trait expression. In both models derived from traditional multivariate quantitative genetics settings (the multilinear model and the gene pleiotropy model), the evolution of mutational correlations was similar: the mutation covariance matrix aligned with the direction of the fitness function, and correlations remained modest. In contrast, in gene regulatory networks, correlations could also evolve qualitatively, but the response was not isotropic in the phenotypic space. Correlation patterns were mostly driven by the regulatory distance and nature (activation or inhibition) of regulations between genes. Simulations confirmed that the correlation between gene expressions decreased with the network distance among genes, and that strongest correlations were associated with direct regulations. As correlated stabilizing selection promotes mutational correlation, it thus promotes network topologies where selected genes are closely connected. Taken together, our results exemplify how the shape of a constant, stabilizing fitness function can theoretically drive the structure of the underlying genetic architecture.

From a multivariate quantitative genetics point of view, organizing the phenotypic space along measurable traits is a necessary consequence of how phenotypes are estimated empirically, but most models can easily be redefined for any linear combination of traits. For instance, when the bivariate multilinear model is parameterized in such a way that epistasis and pleiotropy are identical in all directions of the phenotypic space, mutational effects evolve indifferently towards robustness or pleiotropy depending on the direction of the fitness function (Jones *et al*., 2014 and our simulations). This is where the quantitative genetics predictions break when applied to gene networks: in network models, genes both define observable phenotypes and structure the regulatory patterns. As a consequence, correlated selection can also induce the evolution of pleiotropic gene expression, as predicted by theory, but the mechanisms involved into this response are different from those involved into the evolution of robustness. Our simulations pointed out the major effect of the network distance on co-expression, showing that evolving pleiotropic gene expression requires to rewire the network and reduce the distance (and possibly the sign of regulations) between co-selected genes.

### Model properties

All three models implemented in our simulation software are based on fundamentally different principles, although they all allow for the evolution of pleiotropy.

In the multilinear model, pleiotropy arises as a consequence of the gene-gene interactions (epistasis). Under stabilizing selection, the genetic contributions (*y*^*i*^ in the mathematical model description) can evolve at all genes, provided that the changes are compensated by other genes (so that the phenotype remains constant). Due to the non-linear genotype-phenotype mapping, the average effect of a mutation can thus evolve when the genetic background changes, which modified mutational variances and covariances.

The interaction coefficients *ε* have an empirical meaning, as they measure the curvature of the genotype-phenotype map in some specific directions (Hansen and Wagner, 2001; Le Rouzic, 2014). Yet, the number of coefficients grow very fast with the complexity of the model (proportionally to *n*^*o*^*K*^3^ for a *n*-gene *K*-trait model when considering epistatic interactions of order *o*), and even our simple setting (*n* = 6, *o* = 2, *K* = 2) made it complicated to control the properties of the genetic architecture by finely setting these coefficients. In particular, isotropy (the fact that traits were interchangeable) is not a general property of the multilinear model, but rather a consequence of distributing *ε* coefficients independently.

In contrast, the gene pleiotropy model is perfectly additive, and the effects of mutations are constant throughout the simulations. Each gene has a specific, non-evolvable pleiotropic pattern, and the mutational covariances change at the genotype level because the differential mutation rates of all genes can evolve. It is thus the relative gene contribution to the **M** matrix that drives the evolution of mutational correlations. Contrary to the multilinear model, changing the size of the **M** matrix was not possible (i.e., robustness to mutations could not evolve for both traits at once), and only the shape and the direction of the mutational covariance matrix was evolvable.

The gene network model used for the simulation was based on the popular ‘Wagner’ model (Wagner, 1994, 1996). This model has already been explored in evolutionary biology to study the evolvability, the modularity, and the canalization of gene regulatory networks (Siegal and Bergman, 2002; Ciliberti *et al*., 2007; Rünneburger and Le Rouzic, 2016, see Fierst and Phillips, 2015 for review). This model is computationally fast and requires few parameters in addition to the structure of the regulation network itself. Contrary to both previous models, which were based on quantitative genetics (statistical) principles, the gene network model implements mechanistic interactions among genes, so that complexity emerges from the model structure. More specifically, epistasis emerges from the non-linearity of the sigmoid regulation scaling function, and pleiotropy is due to the causal relationship between the expression level of regulatory genes and the consequences on the expression of regulated genes. Its lack of realism at the biochemical and cellular level (discrete time steps, no degradation kinetics, no compartments, arbitrary dose-response function) makes it less popular for physiological models of known regulation pathways (alternative models could be found in e.g. Karlebach and Shamir, 2008), but it is a convenient framework for theoretical studies of network evolution on evolutionary time scales.

Theoretical approaches to the evolution of genetic architectures are often limited by the oversimplification of the selective constraints. In a multicellular organism, phenotype is many-dimensional and encompass morphological, behavioral, and physiological traits in a complex temporal and spatial context, accounting to different developmental stages, different cell types, and different environmental conditions. In contrast, the Genotype-to-Fitness map in our simulations was simple (multivariate bell-shaped) and phenotypic optima were constant and close to their initial value (no adaptive evolution in the simulations). Realistic patterns and strengths of selection in high phenotypic dimensions are not really known, but recent statistical and experimental progress makes it possible to expect reliable empirically-based estimates in the near future, e.g., for gene expressions in a transcriptome (Whitehead and Crawford, 2006; Koch and Guillaume, 2020; Price *et al*., 2022).

### Co-expression in gene regulatory network

Whether or not stabilizing selection could affect gene network topology is not a trivial question. It is not clear whether structural features of biological networks result from an adaptive process. For instance, modularity can emerge from different mechanisms, including direct selection for efficiency (Clune *et al*., 2013), adaptation to modular environments (Kashtan and Alon, 2005), indirect selection for evolvability, or mutation bias (Wagner *et al*., 2007). Large-scale mathematical properties, such as scale-freeness, may not have any impact on fitness, and the evolution of gene networks may be dominated by non-adaptive mechanisms, including genetic drift and mutation bias (Lynch, 2007). The mechanisms of gene regulation generate a lot of epistasis and pleiotropy at the gene expression level, but these are not expected to be uniformly distributed in the phenotypic space. In particular, coexpression among the genes belonging to the same regulatory module is unavoidable, suggesting that evolving expression independence might be more difficult than evolving correlated expression, as it requires to rewire the network and change its modularity.

We observed repeatedly that gene networks were evolving to match the fitness function qualitatively, but often failed to align to the correct direction. We could discard the possibility that some networks could be trapped at a local optimum, since starting close to the direction of the fitness function evolved to imperfect alignment. Several hypotheses can be proposed to explain this gene-network specific observation : (i) mutations affecting gene co-expression have direct negative side effects (change in gene expression, decrease of the network stability), so that the fitness peak corresponds to a sub-optimal pleiotropic pattern; (ii) mutations affecting gene co-expression have indirect negative side effects (e.g., increase the size of **M**) and actually do not decrease the genetic load; (iii) some **M** matrices cannot be obtained with this gene network model. It was difficult to investigate this question further based on our simulation setting.

Here, we showed theoretically how correlated stabilizing selection on gene expression could deterministically alter the topology of gene networks, by shortening the network distance between genes which expression levels interact at the fitness level. Simulations show that the evolution of regulatory connections as a result of correlated selection can realistically happen in non-restrictive conditions, even if it is difficult to estimate the extent by which real gene networks are affected by this phenomenon. Nevertheless, correlated selection is not the only form of selection that may promote specific network topologies. Directional selection on a multivariate phenotype may for instance indirectly favor genetic backgrounds generating mutational variation towards the optimum. Fluctuating selection could similarly promote pleiotropy when the optimal phenotypes are correlated among traits (Crombach and Hogeweg, 2008).

Independently, fluctuating selection also opens the possibility for the organisms to gather cues about the environment and evolve an adaptive plastic response (Via and Lande, 1985). If, as intuited by Waddington (1942), complex genetic architectures respond to mutational and environmental disturbances through shared molecular mechanisms, it is likely that mutational and environmentally-induced co-expressions will be similar – a property of complex genetic system that could fasten genetic adaptation (Brun-Usan *et al*., 2021; Chevin *et al*., 2021). Direct selection for correlated gene expression plasticity among sets of genes thus appears as a potentially powerful force that could drive the evolution of the topology of gene networks, perhaps strong enough to overcome the influence of correlated selection illustrated in our simulations.

Even if our results explored the theoretical possibility for selection to affect the modularity of genetic architectures, interpreting gene network topologies as systematic consequences of an adaptive process would be largely premature. The influence of correlated selection on the topology and on co-expression in real gene networks remains virtually unknown. Adaptive (e.g., plasticity) and non-adaptive (mutation bias or genetic drift) forces are also prone to alter network topology, and condition the long-term evolvability of these complex genetic architectures. Since gene networks can be shaped by selection, but can also constrain the mutational availability of evolutionary path, the long-term influence of past environment on phenotypic variability and evolvability remains a challenging question in both quantitative genetics and evolutionary systems biology.

## Acknowledgments

We thank Anne Genissel for valuable advice and discussion. Simulations were performed on the Core Cluster of the Institut Français de Bioinformatique (IFB) (ANR-11-INBS-0013).

## Supplementary material

### Supplementary Methods

#### Mutational direction

The mutational direction *α*(**M**) is the angle between trait *a* and the main axis of the 2 *×* 2 mutational matrix **M** between the two focal traits *a* and *b*. It was expressed in the interval (−*π*/2, *π*/2), and calculated as:

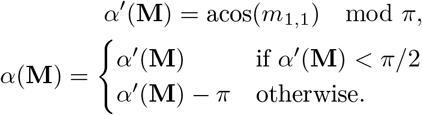

where *m*_1,1_ is the first element of the first eigenvector of **M**.

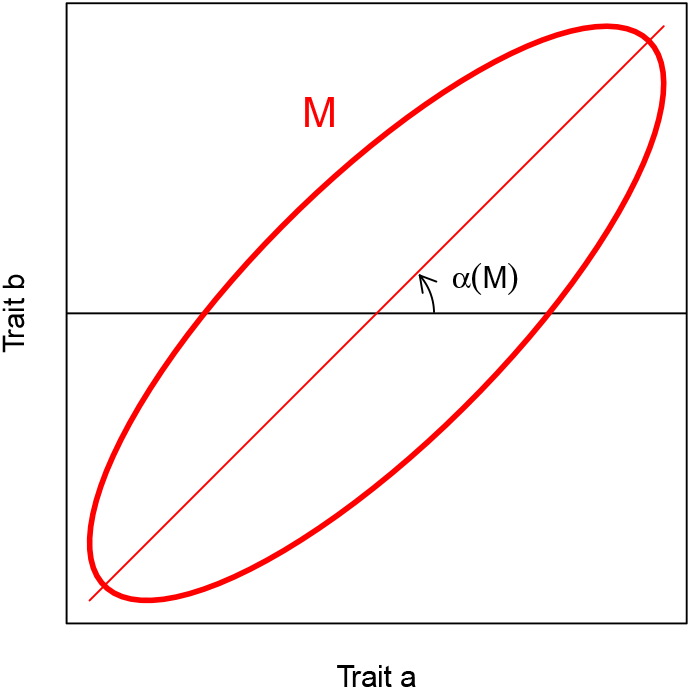

Whenever necessary, the mean direction 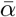 over *R* replicates was obtained by a circular mean restricted to the (−*π*/2, *π*/2) interval:

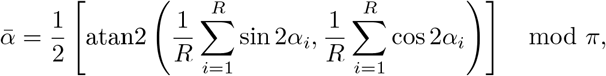

where atan2(*x, y*) = 2 atan 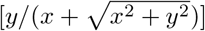 is the angle between the X-axis and the vector (*x, y*).

#### Regulation scaling function

Quantitative gene network models require a scaling function that maps the strength of regulation on a gene (the total effects of all transcription factors acting on the gene) and gene expression. Here, we used the same scaling function as in Rünneburger and Le Rouzic 2016:

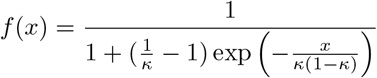

where *κ* ∈ (0, 1) stands for the basal expression level. By definition, *f* (0) = *κ* (in absence of regulation, the gene is expressed at its basal level), and the function is scaled so that d*f*/d*x* |_*x*=0_ = 1, in order to ensure that effects of genotype changes are comparable across simulations with different basal levels.

With the default basal expression *kappa* = 0.5, the scaling function reduces to *f* (*x*) = 1/(1 + *e*^−4*x*^).

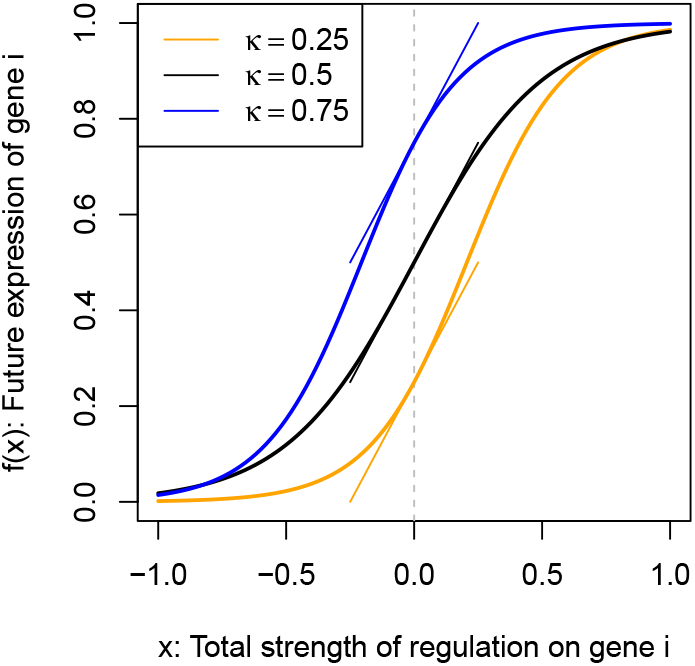

In the simulations, the regulation on gene *i* was obtained by adding up the effect of transcription factors, proportionally to their concentration: *x*_*i*_ = ∑_*j*≠*i*_ *P*_*j*_*W*_*ij*_.

#### Selection against unstable networks

The fitness component associated with selection against unstable (cyclic) networks was a negative exponential function of the variance in gene expression. This exponential scaling ensures that the fitness penalty is nil when the network is stable (*w*_stab_ = 1 when ∑*V*_*i*_ = 0), and that the individual is virtually not viable when the network is unstable (*w*_stab_ → 0 when ∑ *V*_*i*_ is large).

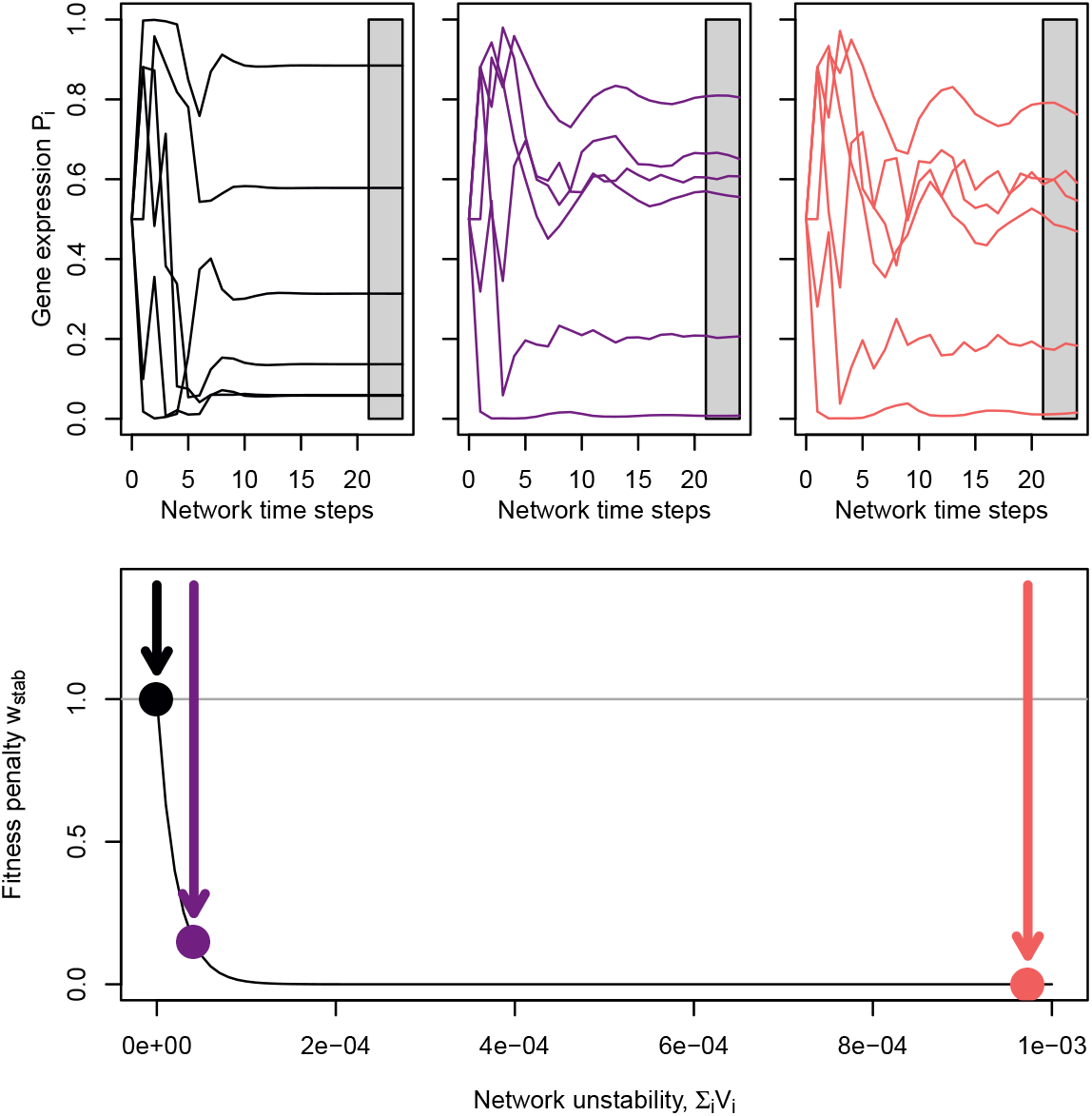

The figure displays the dynamics of three arbitrary 6-gene networks displaying diverse stability behavior. Gene expression stability is measured in the gray areas (steps 21 to 24). The bottom panel represents the fitness penalty used in the simulations (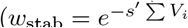, with *s*^*′*^ = 46, 000). The fitness effect associated with the stable network (black, left) is *w*_s*tab*_ ≃ 1, which does not penalize the fitness function. In contrast, fitness is multiplied by *w*_s*tab*_ ≃ 0 for the unstable network (red, right), making the individual unviable regardless of the gene expression level. With the strong selection coefficient *s*^*′*^, even slightly fluctuating networks (violet) were substantially penalized.

## Supplementary Figures

**Supplementary Figure 1:**
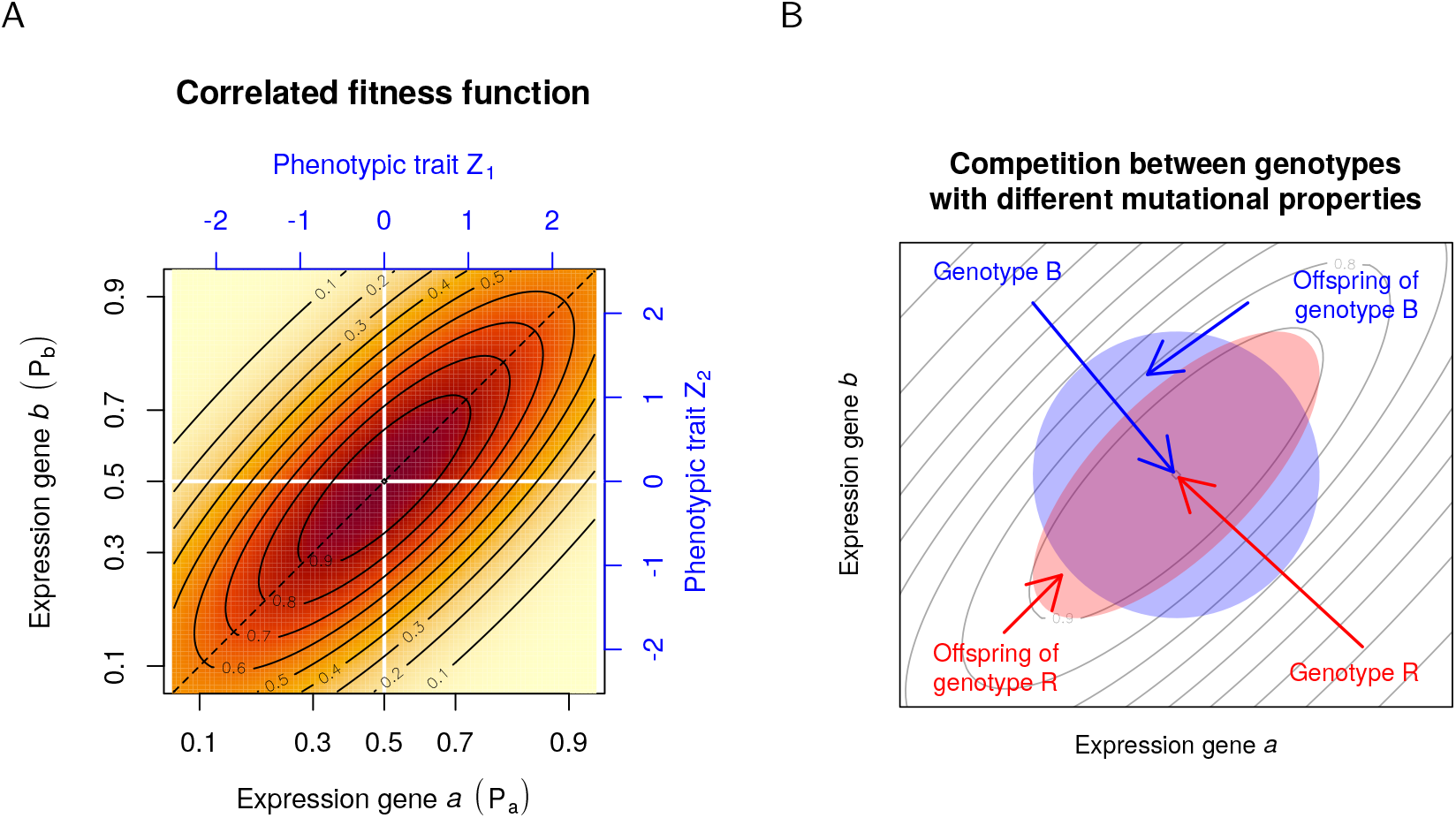
A: representation of the bivariate correlated fitness function used in the simulations (here, with an direction *α*(**M**) = *π*/4). Axes display both gene expression levels (**P**) scaled between 0 (no expression) and 1 (full expression) in black, and the corresponding rescaled phenotypes **Z** from which fitness was computed in all three models (in blue). White lines highlight the optimal phenotype (for which the fitness is maximal). Any deviation from the optimal phenotype is penalized, but the penalty is weaker when both traits change together. B: Cartoon representation of the advantage of a genotype in which mutational effects are correlated (red, R) over a genotype in which mutational effects are uncorrelated (blue, B). Both genotypes display the optimal gene expression and thus have the same fitness; mutant offspring from both genotypes deviate from the optimum within the same range, but due to the genetic correlation, the average fitness of the mutant offspring (iso-fitness lines in gray) from genotype R is higher than the offspring from genotype B: the R lineage will progressively replace the B lineage in the population.

**Supplementary Figure 2:**
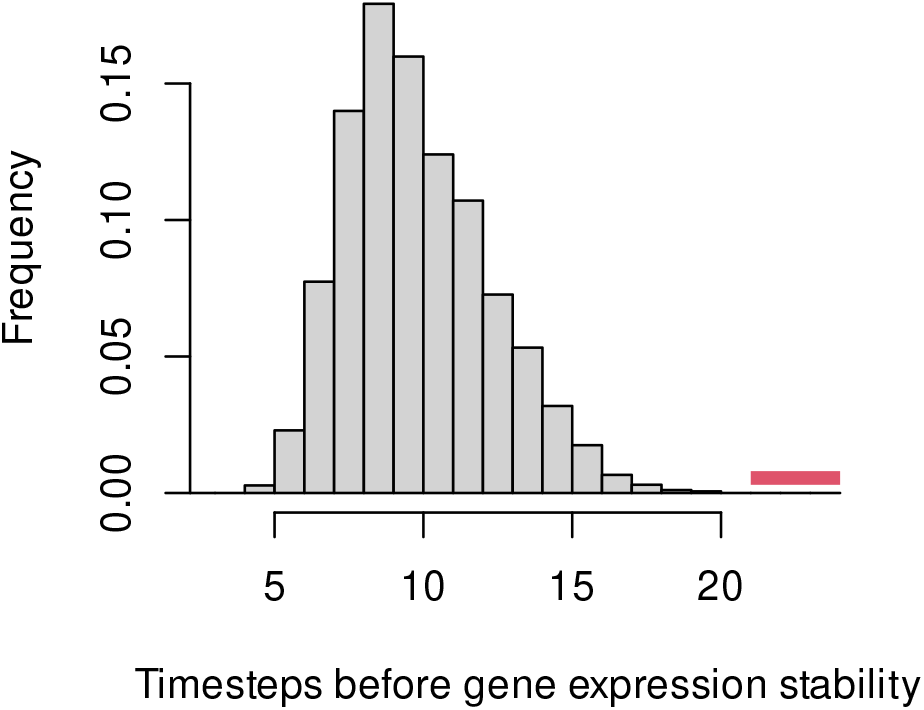
Number of dynamic steps during the “development phase” necessary for genes to reach a stable expression, i.e a variance between time-steps *<* 0.0001. The equilibrium expression is the mean expression of time-steps 21 to 24, indicated in red. The last generation of all of our 8281 GRN simulations presented in the main text are represented in this figure.

**Supplementary Figure 3:**
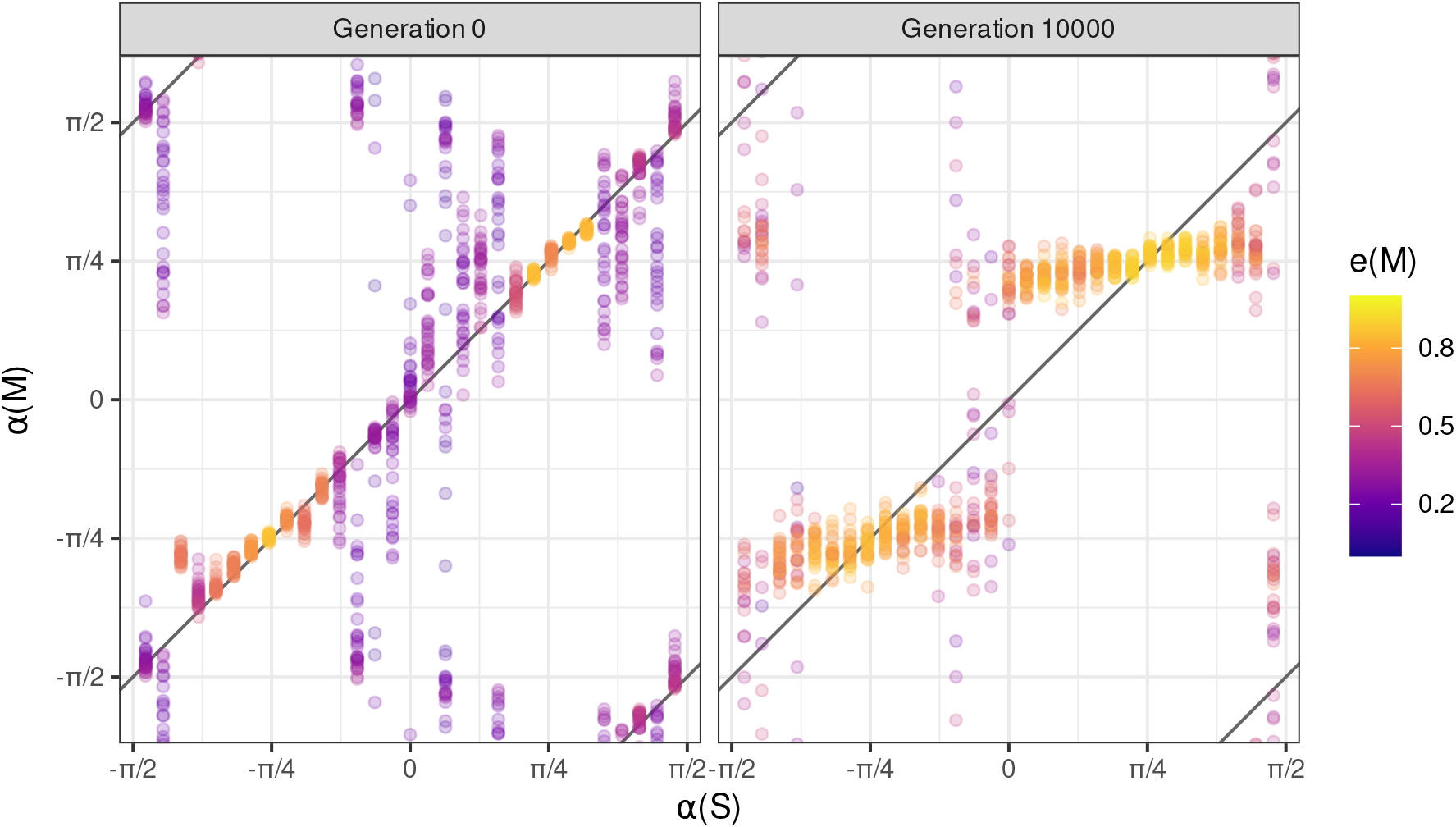
Evolution of *α*(**M**) in response to *α*(**S**), starting simulations with networks giving the closest alignment in previous simulations (Figure 2C, GRN model). For the same *α*(**S**), replicates started with the same GRN. The variance obtained in the *α*(**M**) at generation 0 is due to sampling effects when computing **M**, which was substantial when **M** were close to round (no well-defined direction).

**Supplementary Figure 4:**
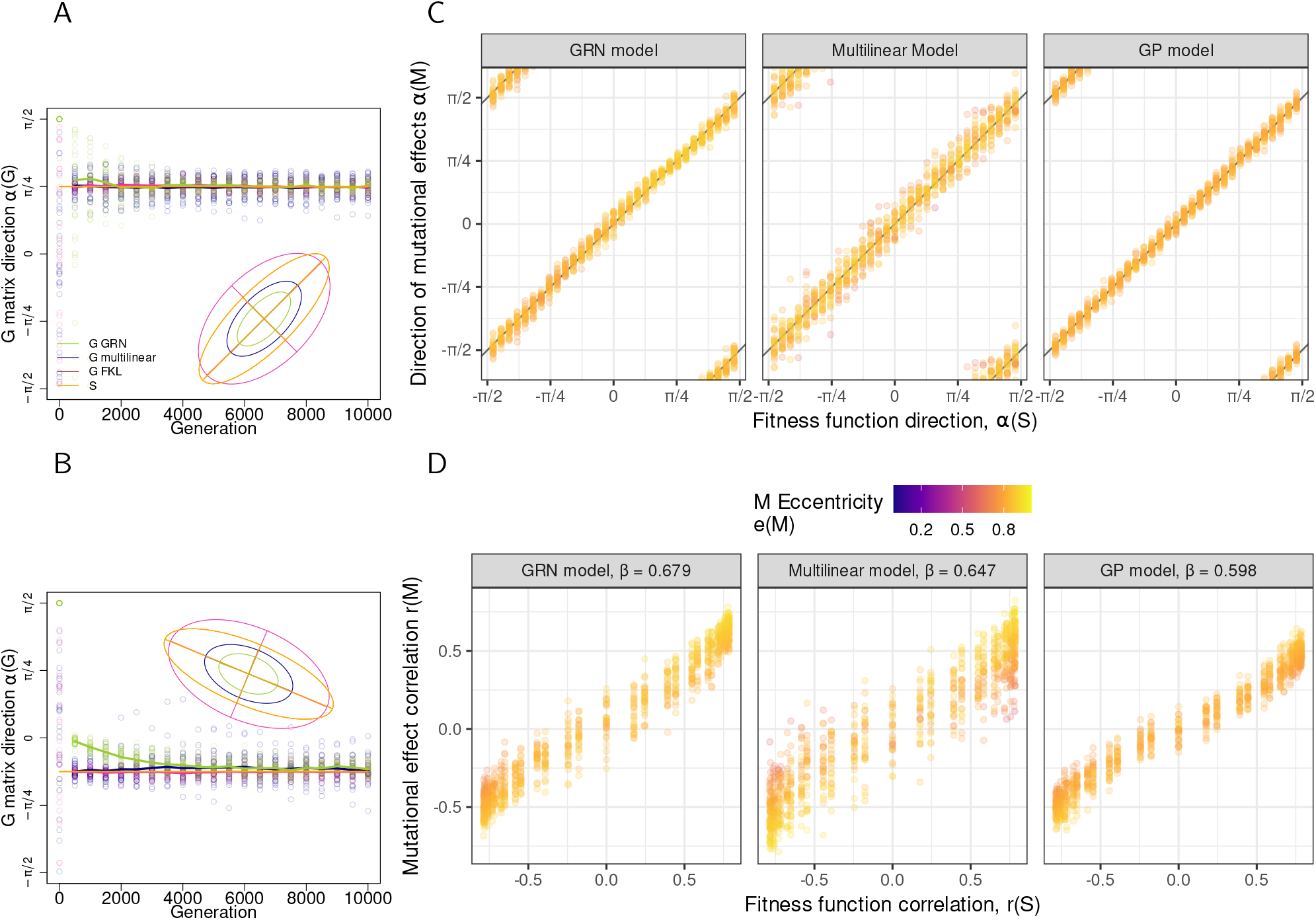
The same as Figures 2, but for the **G** matrix. A, B: Evolution of the angle of the main axis of the mutation matrix (*α*(**G**)) along generations. Orange ellipses represent the fitness function (scaled × 0.025), which direction was *α*(**S**) = +*π*/4 (panel A), and *α*(**S**) = − *π*/8 (panel B). Dots illustrate 30 simulation replicates, plain lines stand for circular means 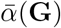. Differences in the shape of **M** and **G** matrices can be explained by linkage disequilibrium (the more LD, the more **G** can be similar to **S**). In A and B, the sizes of the **G** matrix differ between the models more than their **M** matrix, due to to different responses of the three models to LD. The **G** matrix in the GP model is the least affected by linkage disequilibrium, as evolution tends to decrease the effective number of loci contributing to the traits. In contrast, regulatory sites in the promoter of genes are completely linked in the GRN model, allowing for the evolution of strong and persistent LD.

**Supplementary Figure 5:**
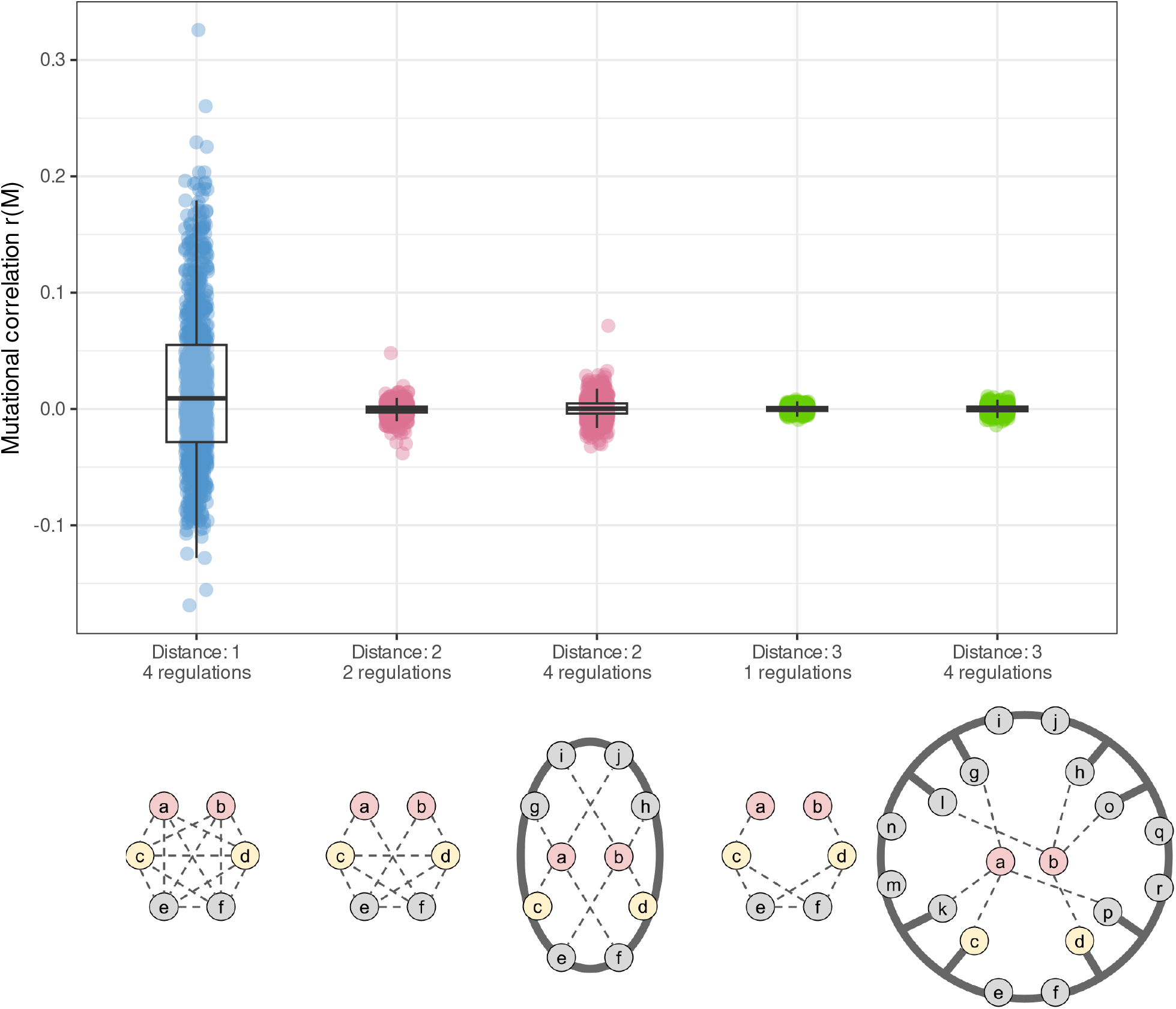
Distribution of mutational correlations with different networks (same color code as in Figure 5). Two topologies were compared for the same network distance : one with the number of genes conserved (6 genes), and one with the number of edges connected to *a* and *b* conserved (4 possible regulations). Genes located on grey circle are all connected to each other; genes connected to the grey circle can interact with every gene on the circle but not with each other.

**Supplementary Figure 6:**
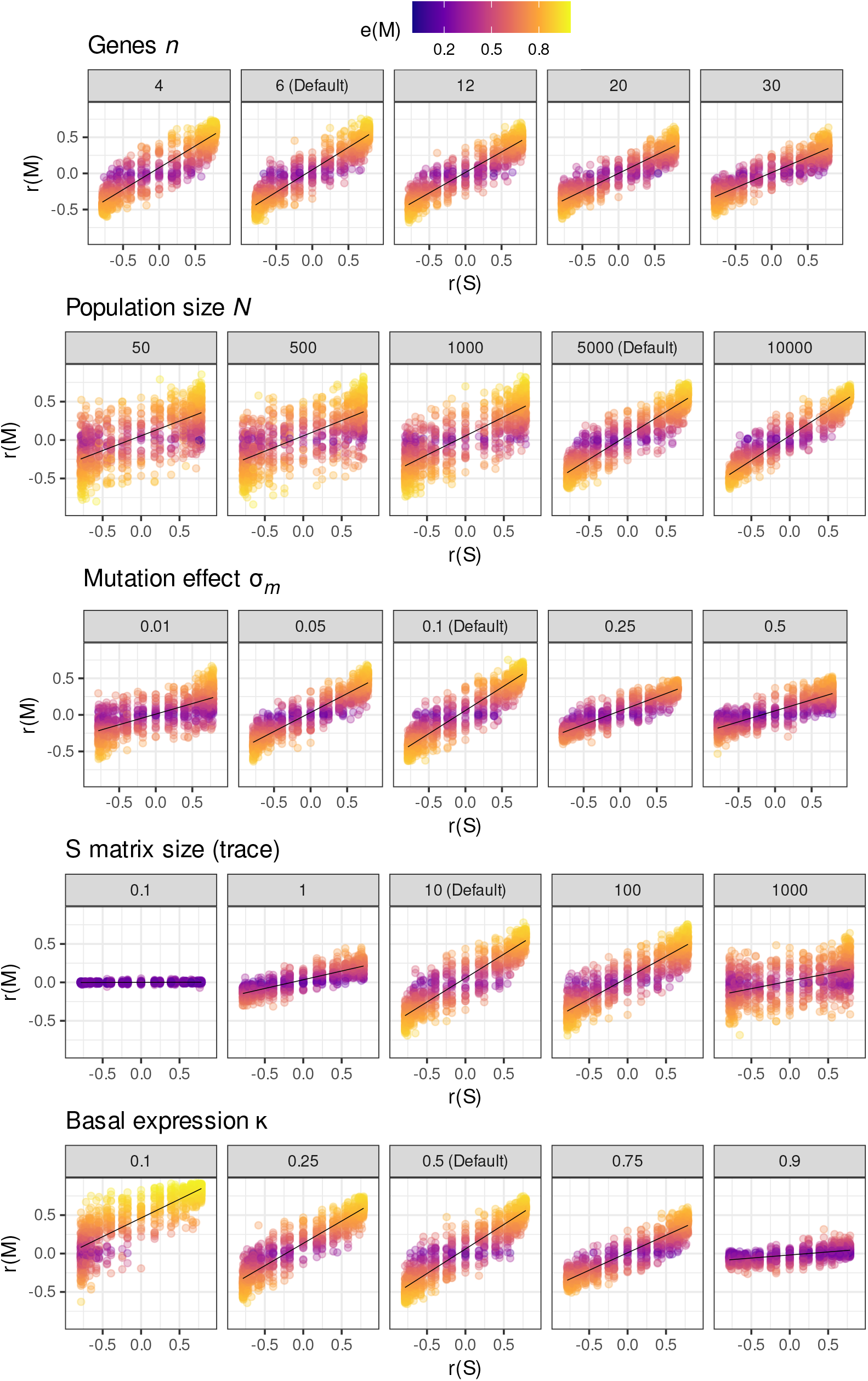
Parameter exploration on the response from the mutational correlation *r*(**M**) to the selection correlation *r*(**S**). When changing the network size *n*, the mutation rate per individual *μ* was adjusted to keep the same mutation rate per gene. For the smallest values of *σ*_*m*_ 0.01 and 0.05), the simulation duration was changed to 100000 and 20000 generations, respectively, to ensure that the population has reached a similar equilibrium. Gene expression optima ***θ*** were adjusted to follow the basal expression *κ*.

